# Microbiome-Targeted Reduction of Circulating Trimethylamine N-Oxide Mitigates Ischemic Stroke Risk

**DOI:** 10.64898/2026.04.15.718846

**Authors:** Jin Sun You, Chung Eun Yoon, June Beom Kim, Mohammed Abd Alrahman, Hee Yong Jung, Mi Young Yoon, You Bin Kim, Sang-Guk Lee, Hyo Suk Nam, Sang Sun Yoon

## Abstract

Elevated plasma trimethylamine N-oxide (TMAO) is an independent predictor of major adverse cardiovascular events and ischemic stroke. While inhibition of microbial TMA production has been explored, concerns regarding off-target effects and limited efficacy in complex microbial ecosystems have hindered clinical translation. Here, we report a microbiome-based therapeutic strategy based on the direct enzymatic degradation of intestinal TMA by *Paracoccus aminovorans* BM109. Through targeted screening, we identified BM109 as a commensal strain harboring a comprehensive set of enzymes capable of metabolizing TMA and TMAO into non-toxic end products under both aerobic and anaerobic conditions. In a chronic high-choline diet murine model, oral administration of BM109 resulted in a 38% reduction in systemic TMAO levels. In a rat model of transient middle cerebral artery occlusion (tMCAO), short-term pre-treatment reduced cerebral infarct size by 58% and significantly improved neurological outcomes. These effects were accompanied by favorable safety observations, including the absence of hemolytic activity and intestinal tissue damage. Collectively, our findings establish BM109 as a promising live biotherapeutic product that targets the gut microbiome-host metabolic axis. By reducing the systemic TMAO burden, BM109 represents a potential strategy for modulating cardiometabolic and cerebrovascular risk.

## Introduction

The human body maintains its function and health through intricate metabolic processes and reciprocal interactions with commensal microbes, collectively termed the microbiome (*1, 2*). Studies estimate that approximately 40 trillion microbial cells inhabit the human body (*3*), outnumbering human cells in a ratio of approximately 1.3:1 (*4*). This vast microbial community profoundly influences human health, functioning as an integral metabolic organ. Advances in microbiome research have highlighted the role of the gut microbiome in a wide range of systemic diseases beyond the gastrointestinal tract, including metabolic disorders (*5*), neurodegenerative conditions (*6*), kidney dysfunction (*7*) and cardiovascular diseases (CVDs) (*8*).

Among the diverse functions of the gut microbiome, its role in cardiovascular diseases has gained increasing attention, particularly in conditions such as stroke (*9*), atherosclerosis (*10*) and thrombosis (*11*). Beneficial gut microbes contribute to cardiovascular health by producing short-chain fatty acids (SCFAs) (*12, 13*). SCFAs, particularly butyrate, produced by species such as *Roseburia intestinalis*, exhibit atheroprotective effects, mitigating inflammation and improving vascular function (*14*). In addition, certain gut bacteria exert antioxidant and anti-inflammatory properties, preventing oxidative stress-related endothelial damage (*15, 16*).

Recent studies indicate that gut microbial modification and cardiovascular disease can mutually affect each other (*17*). Metabolomics studies have illustrated that patients with cardiovascular disease display altered composition and diversity of gut microbiome, different from the eubiotic state (*18–20*). In addition, several studies postulate that an increase or decrease in specific bacteria can lead to changes in gut metabolites, such as bile acid or trimethylamine (TMA), which can potentially threaten cardiovascular health (*21–24*). Studies have demonstrated that trimethylamine (TMA), a gut microbiome–derived metabolite, contributes to cardiovascular disease through its hepatic conversion to trimethylamine N-oxide (TMAO) by flavin-containing monooxygenase 3 (FMO3) (*25*). TMAO promotes atherosclerosis and thrombosis by enhancing vascular inflammation, lesion formation, immune cell activation, and platelet reactivity (*23, 26*). Mechanistically, TMAO exacerbates endothelial dysfunction by triggering mitochondrial ROS production and inhibiting the Nrf2-mediated antioxidant defense system, leading to vascular inflammation and pyroptosis (*27*). Clinically, elevated circulating TMAO levels are consistently associated with increased cardiovascular risk and adverse outcomes, and have been identified as an independent predictor of major adverse cardiovascular events (MACE) across multiple cohort studies (*28–34*). Elevated TMAO levels pose additional health risks beyond cardiovascular pathology. In patients with chronic kidney disease (CKD), impaired renal clearance results in persistently high plasma TMAO levels, exacerbating cardiovascular complications, one of the leading causes of mortality in this population (*7, 35, 36*). Despite the strong and consistent associations observed in epidemiological studies, interventional approaches aimed at lowering circulating TMAO levels remain limited.

A major challenge in targeting gut-derived TMA and TMAO lies in their continuous production within the gut environment, largely independent of microbiome composition. A wide range of bacterial taxa, including members of the genera Clostridium, Shigella, Klebsiella, and Citrobacter, are capable of metabolizing dietary substrates such as choline, betaine, carnitine and related compounds to generate TMA (*37, 38*). Given that TMA-producing microbes are taxonomically diverse and widely distributed within the gut, selective eradication is unfeasible. Additionally, dietary restriction of TMA precursors such as choline, betaine, lecithin, and carnitine is impractical, as these nutrients play important roles in human physiology.

Since direct suppression of TMA biosynthesis remains challenging, an alternative strategy is to degrade TMA post-production to prevent its systemic absorption. In this study, we sought to identify commensal gut bacteria with intrinsic TMA-degrading capabilities. Through targeted screening of human fecal microbiota, we isolated a commensal species exhibiting robust enzymatic activity against TMA and TMAO. We characterized its metabolic pathways, assessed its efficacy in reducing TMAO levels under aerobic and anaerobic conditions, and validated its *in vivo* therapeutic potential using both a diet-induced and a disease-relevant animal model. This study provides a proof-of-concept for a microbiome-based TMAO-lowering therapy through direct degradation of microbial TMA.

## Results

### Isolation and Identification of a Robust TMA-Degrading Bacterial Species from Human Feces

Methylotrophic bacteria are known to utilize C1 compounds such as methane and methanol, and can also metabolize trimethylamine (TMA) and trimethylamine N-oxide (TMAO), which contain three methyl groups (*39*). Based on this metabolic feature, we sought to isolate fecal bacteria with TMA-degrading capacity.

As outlined in Fig. 1A, pooled human fecal samples were screened under conditions favoring methylotrophic growth, yielding multiple candidate colonies. These isolates were subsequently evaluated for their ability to degrade TMA. Among the screened candidates, several strains exhibited measurable TMA consumption, with a subset demonstrating robust degradation activity after 24 h incubation (Fig. 1B). These strains were selected for further characterization, and TMA levels were quantified using an optimized picric acid-based assay (*40*), as described in Methods.

**Figure 1.**
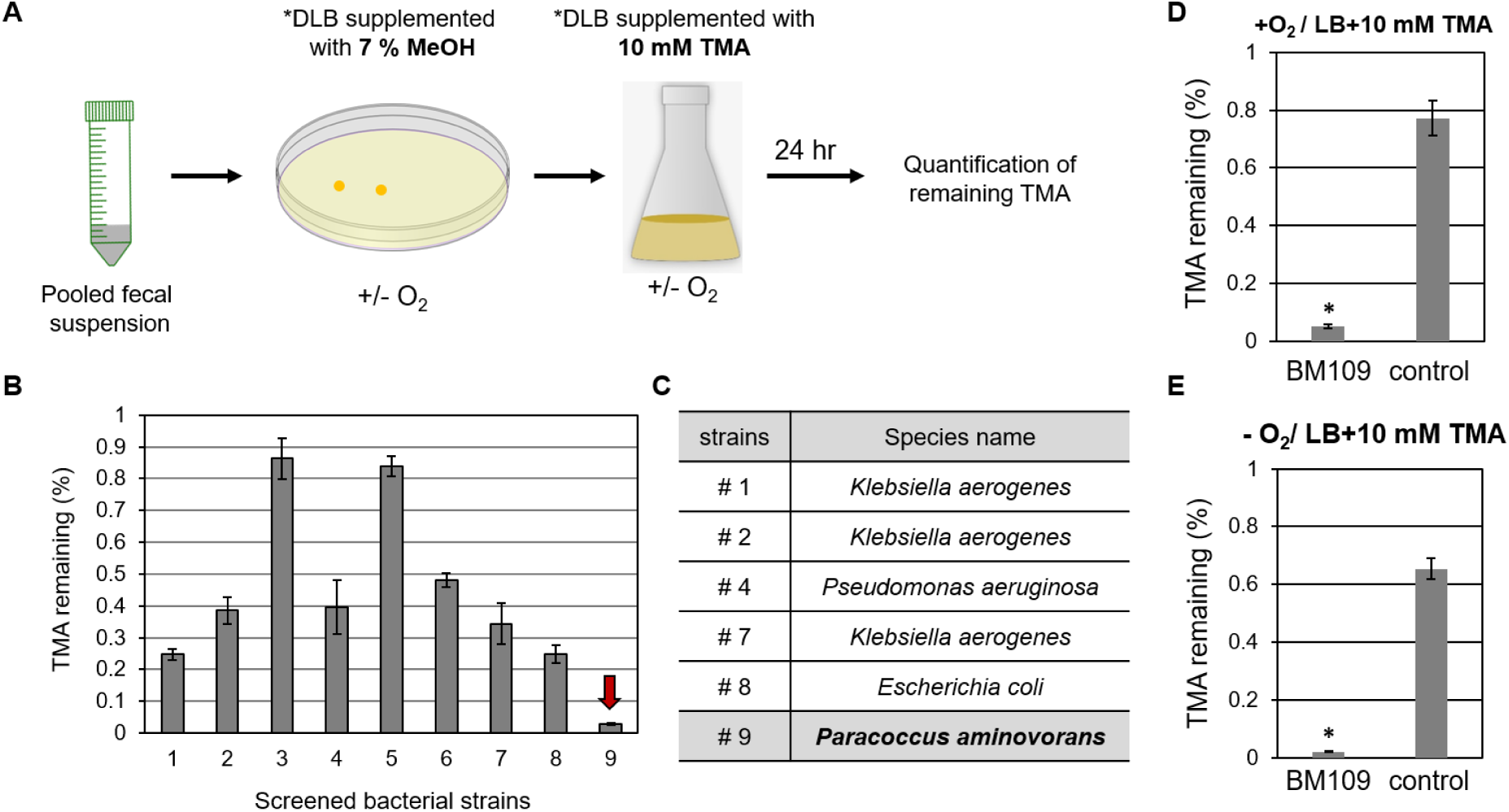
Schematic procedure of screening for TMA-degrading fecal microbes. (**A**) Schematic representation of the screening procedure. Human fecal samples were pooled to generate a microbiome suspension, which was plated onto DLB agar supplemented with 7% MeOH to enrich for methylotrophic bacteria. After 48 h of incubation, isolated colonies were inoculated into liquid DLB medium supplemented with 10 mM TMA to assess their growth and TMA-degrading capacity. After 24 h of cultivation, culture supernatants were collected for TMA quantification using a picric acid-based colorimetric assay. *DLB refers to 1/5 diluted LB medium. (**B**) Quantification of residual TMA in culture supernatants after 24 h of aerobic growth in DLB supplemented with 10 mM TMA. The amount of remaining TMA was compared to the initial input and expressed as a ratio (mean ± SD, n = 3). Strain No. 9 exhibited the highest TMA degradation efficiency (red arrow). (**C**) Bacterial strains capable of degrading >60% of the initial TMA were selected for species identification via 16S rRNA gene sequencing. Strain No. 9 was identified as *Paracoccus aminovorans*, while the other strains belonged to *Klebsiella aerogenes*, *Pseudomonas aeruginosa*, and *Escherichia coli*. BM109 and a control strain (*P. denitrificans*) were grown in LB amended with 10 mM TMA aerobically with shaking for 24 h (**D**) and anaerobically without shaking for 48 h (**E**). To support anaerobic respiratory growth, 60 mM NO ^−^ was added in the anaerobic culture media. Concentrations of TMA detected in the supernatants were compared with the initially added TMA and presented with ratio values (mean ± SD, n=3). \**p* < 0.001 versus values obtained from control growth. Under both conditions, BM109 exhibited significantly greater TMA degradation compared to the control strain, while bacterial growth was comparable between the two strains.

Nine strains were selected as primary candidates based on their consistent growth on MeOH-containing plates. Their TMA-degrading capacities were evaluated by quantifying residual TMA levels relative to the initial concentration. Among these, seven strains degraded more than 50% of TMA, although their activities varied (Fig. 1B). Notably, strain 9 exhibited near-complete TMA degradation within 24 h under aerobic conditions. In contrast, strains 3 and 5 showed minimal TMA degradation despite robust colony formation, indicating that growth under methylotrophic conditions does not necessarily correlate with TMA-degrading activity in liquid culture.

Species identification based on 16S rRNA gene sequencing revealed that strains 1, 2, and 7 were *Klebsiella aerogenes*, strain 8 was *Escherichia coli*, and strain 4 was *Pseudomonas aeruginosa* (Fig. 1C), all of which are known opportunistic pathogens (*41, 42*). In contrast, strain 9 (hereafter referred to as BM109) was identified as *Paracoccus aminovorans* (99.91% sequence identity to JCM7865), a member of the alpha-Proteobacteria distinct from the other isolates (*43*). Notably, Paracoccus species have been reported to be present in the human gastrointestinal tract (*44, 45*).

To evaluate TMA-degrading efficacy under gut-relevant conditions, BM109 was cultured in LB medium and assessed under both aerobic and anaerobic conditions. BM109 degraded∼95% of TMA within 24 h under aerobic conditions (Fig. 1D). Under anaerobic conditions with nitrate as an alternative electron acceptor, BM109 degraded ∼97.9% of TMA within 48 h (Fig. 1E), despite reduced growth. These results demonstrate robust TMA-degrading activity across oxygen conditions, supporting its potential functionality in the gut environment. As a control, *Paracoccus denitrificans*, a related species within the Paracoccus genus, exhibited markedly lower TMA-degrading activity, highlighting the distinct metabolic capability of BM109.

### Genomic Structure and Comparative Analysis of BM109

The entire genome of BM109 was sequenced and assembled into four contigs, two of which contained both tRNA and rRNA gene sequences. The sequencing generated a total of 14,415,666 reads with an average depth of 473.9, and the total genome size was determined to be 4,297,217 bp. Genome annotation revealed 4,095 coding sequences (CDS) across all four contigs, with detailed features summarized in Table 1. The genome sequence data have been deposited in the NCBI BioProject database under accession number PRJNA891425.

**Table 1.** Overall landscape of BM109 whole genome and comparison with *P. aminovorans* JCM7685 genome. BM109 genome is constituted in 4 different contigs, with the first contig being the largest chromosome 1. Overall genome architecture is similar to that of JCM7685. GC content (%) represents the total guanine and cytosine ratio in each DNA contig. CDS stands for coding sequence and indicates the number of genes coding for polypeptide products. rRNA and tRNA indicate the number of genes encoding rRNA and tRNA, respectively.

| Strain | type | Size (Mb) | GC (%) | CDS | rRNA | tRNA | contig |
| --- | --- | --- | --- | --- | --- | --- | --- |
| **BM109** | Chromosome I | 2.93 | 67.5 | 2,849 | 6 | 52 | 1 |
|  | Plasmid II | 0.19 | 68.6 | 154 | 0 | 0 | 3 |
|  | Plasmid III | 0.02 | **56.9** | 21 | 0 | 0 | 4 |
|  | Plasmid IV | 1.16 | 66.5 | 1,071 | 3 | 6 | 2 |
|  | **total** | **4.30** | **67.2** | **4,095** | **9** | **58** |  |
| **JCM7685** | Chromosome I | 3.11 | 67.2 | 2,956 | 6 | 47 |  |
|  | Plasmid II | 0.19 | 68.8 | 145 | 0 | 0 |  |
|  | Plasmid III | 0 | 68.4 | 4 | 0 | 0 |  |
|  | Plasmid IV | 0.74 | 67.8 | 620 | 3 | 3 |  |
|  | **total** | **4.04** | **67.4** | **3,725** | **9** | **50** |  |

Comparative genomic analysis revealed high 16S rRNA sequence similarity (99.91%) between BM109 and *Paracoccus aminovorans* JCM7685 (Fig. 1C), but substantial divergence at the whole-genome level. Only 7.47% of contig 1 and limited regions of contig 2 aligned with the reference JCM7685 genome (see Supplementary Information), indicating significant genomic variation. These findings suggest that BM109 represents a distinct strain within the *P. aminovorans* species. The overall genome structure of BM109 was similar to that of JCM7685, consisting of four contigs with contig 1 as the largest. However, notable differences in GC content were observed. In particular, plasmid III of BM109 exhibited a markedly lower GC content (56.9%) compared to the other contigs, whereas JCM7685 showed relatively uniform GC content across its genome. This discrepancy suggests that BM109 may have acquired plasmid III through horizontal gene transfer from an external source.

### Functional Genomics of TMA/TMAO Metabolism

Our genome annotation analysis revealed multiple enzymes encoded in BM109’s genome that are associated with the metabolism of TMA and TMAO, as well as their downstream products. These findings enabled us to construct a functional metabolic pathway for the sequential degradation of TMA and TMAO into non-toxic compounds, including CO_2_ (Fig. 2A).

**Figure 2.**
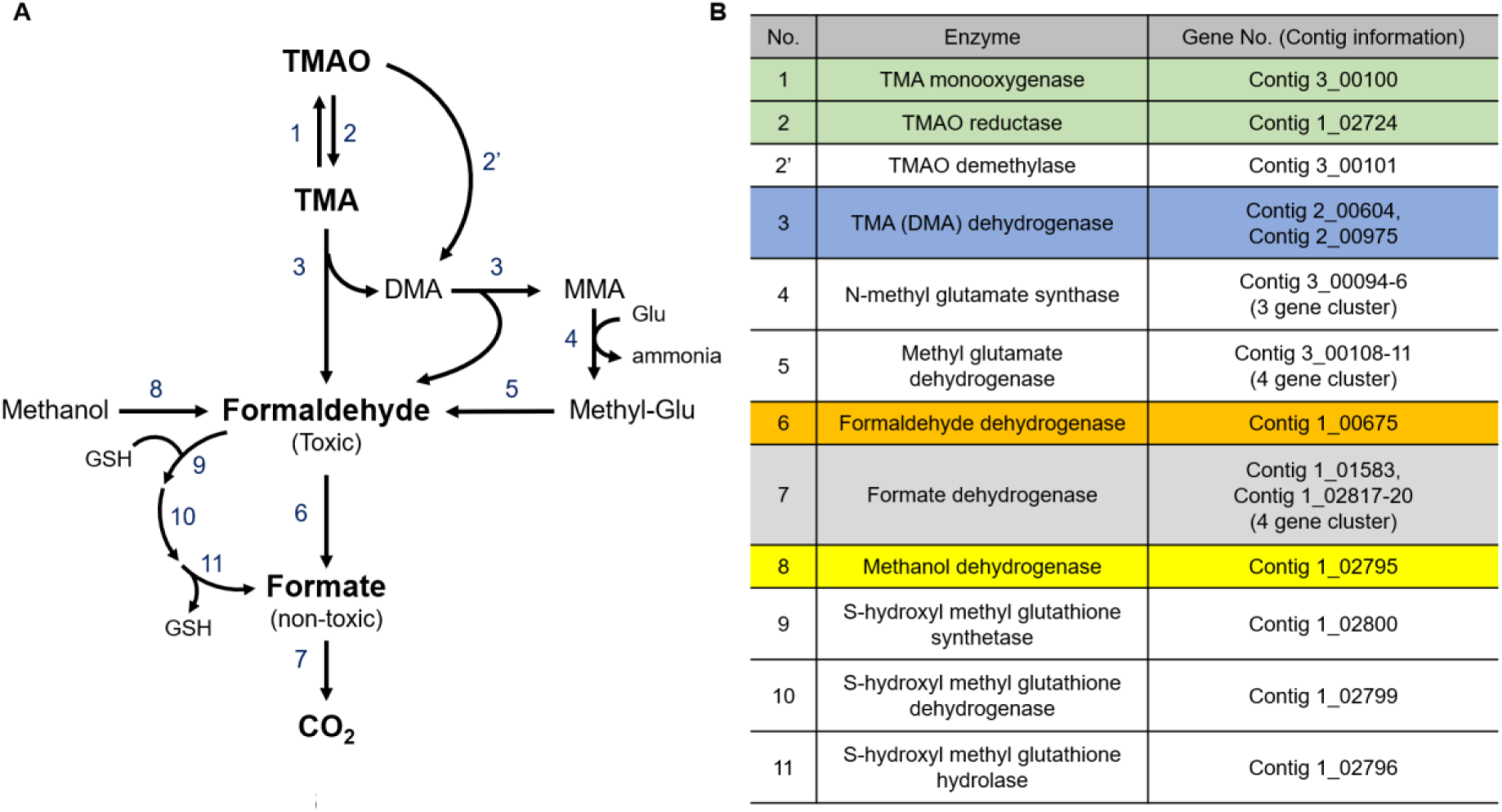
Metabolic pathway of TMA and TMAO degradation and corresponding gene annotations in BM109. (**A**) Schematic representation of TMA and TMAO metabolism. TMA, generated from nutrient digestion, is either oxidized to TMAO by TMA monooxygenase (enzyme 1) or converted into dimethylamine (DMA) and formaldehyde by TMA (DMA) dehydrogenase (enzyme 3). DMA undergoes further metabolism into monomethylamine (MMA) and formaldehyde. Formaldehyde, a toxic intermediate, is processed through multiple detoxification steps. Formaldehyde dehydrogenase (enzyme 6) catalyzes the conversion of formaldehyde into formate, which is subsequently oxidized to CO_2_ by formate dehydrogenase (enzyme 7). Methanol, another potential substrate, can also be converted into formaldehyde by methanol dehydrogenase (enzyme 8). Additionally, TMAO reductase (enzyme 2) enables the reduction of TMAO back to TMA, and TMAO demethylase (enzyme 2’) facilitates alternative TMAO degradation. Detoxification of formaldehyde also involves glutathione (GSH)-dependent pathways, with S-hydroxymethyl glutathione synthetase (enzyme 9), dehydrogenase (enzyme 10), and hydrolase (enzyme 11) contributing to formate production. (**B**) Corresponding genes identified in the BM109 genome encoding enzymes involved in TMA and TMAO metabolism. Each enzyme in panel A is matched with specific gene loci found in BM109, including multiple gene clusters encoding key metabolic enzymes. Notably, two distinct genes (Contig 2_00604 and Contig 2_00975) encode TMA dehydrogenase, and two loci encode formate dehydrogenase, with one organized as a four-gene operon.

Two genes located on contig 2 were annotated as encoding TMA dehydrogenase (TMADH, 2_00604 and 2_00975), an enzyme responsible for converting TMA into dimethylamine (DMA) and formaldehyde (Fig. 2B, highlighted in blue). The presence of two genes encoding the same enzymatic function indicates the importance of genetic redundancy in maintaining critical metabolic functions. The enzymes labeled as “1” and “2” in Fig. 2A correspond to TMA monooxygenase (TMM) and TMAO reductase (TMAORD), respectively (Fig. 2B, highlighted in green). BM109 encodes both enzymes, enabling it to oxidize TMA into TMAO and reduce TMAO back into TMA. This dual functionality allows BM109 to maintain a dynamic TMA/TMAO balance, ensuring metabolic flexibility.

The hydrolysis of TMA produces formaldehyde, a toxic byproduct that must be further metabolized to avoid harm. BM109 contains a gene encoding formaldehyde dehydrogenase (FADH, 1_00675), which catalyzes the conversion of formaldehyde to formate, a less toxic intermediate (Fig. 2B, highlighted in orange). Furthermore, two gene clusters encoding formate dehydrogenase catalyze the oxidation of formate into CO_2_ (Fig. 2B, highlighted in gray). This sequential pathway ensures complete detoxification of formaldehyde, underscoring BM109’s ability to mitigate TMA-derived toxicity.

A gene encoding methanol dehydrogenase (1_02795) was also identified in the BM109 genome, explaining its ability to metabolize methanol to CO_2_ (Fig. 2B, highlighted in yellow). This metabolic feature likely contributed to BM109’s growth under methanol-rich conditions during the initial screening process. Methanol metabolism also produces NADH (*46*), a molecule essential for energy production via respiratory pathways. This capacity likely contributed to BM109’s growth under methanol-rich conditions during the initial screening process.

To evaluate the uniqueness of BM109’s genetic repertoire, we conducted a comparative genomic analysis using BLAST searches. Based on the reconstructed TMAO-to-CO_2_ metabolic pathway (Fig. 2A), we defined a set of 22 proteins encoded by BM109 that are involved in TMA and TMAO metabolism. Notably, this set includes enzymes encoded by multi-gene operons, in which individual gene products were considered separately, resulting in a total of 22 proteins for analysis. The analysis identified 15 Proteobacteria species harboring homologous or similar sequences to these proteins, including nine species within the Alpha-proteobacteria (Fig. 3). However, none of the examined species possessed the complete set of genes required for the full TMAO-to-CO_2_ metabolic pathway. Notably, even *P. aminovorans* JCM7685, the closest relative of BM109, lacked the gene encoding TMAO reductase, which was further confirmed by PCR analysis (Fig. S1).

**Figure 3.**
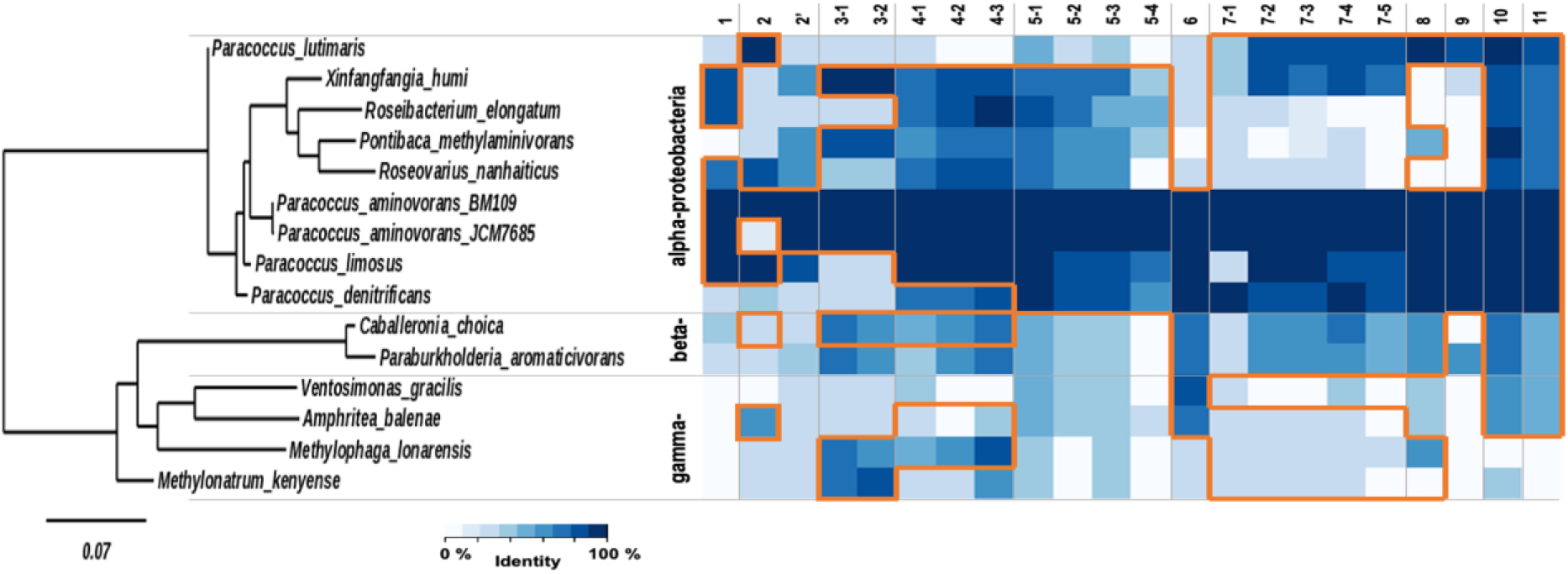
Comparative genomic analysis of BM109 and closely related species based on the presence of TMA and TMAO metabolism-related genes. A total of 15 strains, including two *Paracoccus aminovorans* strains (BM109 and JCM7685), were identified as the closest phylogenetic relatives of BM109, and their phylogenetic relationships were constructed based on 16S rRNA gene sequences. These species were classified into three major groups: nine species from alpha-Proteobacteria, two from beta-Proteobacteria, and four from gamma-Proteobacteria. The heatmap on the right represents the presence and sequence identity of genes associated with TMA and TMAO metabolism across these species. Homologous genes identified via BLAST-based searches using BM109 protein sequences are shown in dark blue (≥60% sequence identity), whereas genes with lower sequence identity but annotated with a similar function are shown in light blue. The four genes within the operon spanning 1_02817∼20 in BM109 are labeled 5b-1, 5b-2, 5b-3, and 5b-4. Genes with conserved presence across multiple species are outlined in orange. This analysis highlights the genomic distinction of BM109 and provides insights into the evolutionary distribution of key metabolic genes.

Among the Alpha-proteobacteria, *P. limosus* exhibited partial coverage of the pathway, encoding more components than most related species, including *P. denitrificans*, but still lacking key enzymes required for complete TMA/TMAO metabolism. Similarly, Beta- and Gamma-proteobacteria species possessed only incomplete subsets of the pathway, missing essential enzymes for full metabolic conversion (Fig. 3).

Collectively, these results demonstrate that BM109 uniquely harbors a complete and functionally integrated genetic repertoire for TMA and TMAO metabolism, distinguishing it from other Proteobacteria species.

### Oral delivery of BM109 reduces TMA and TMAO levels *in vivo*

To evaluate the *in vivo* efficacy of BM109, mice were orally administered BM109 while being fed a normal chow diet (Normal), high-choline diet (HCD), or high-fat diet (HFD) (Fig. 4A). Mice were maintained on these diets for 45 weeks to model long-term dietary exposure.

**Figure 4.**
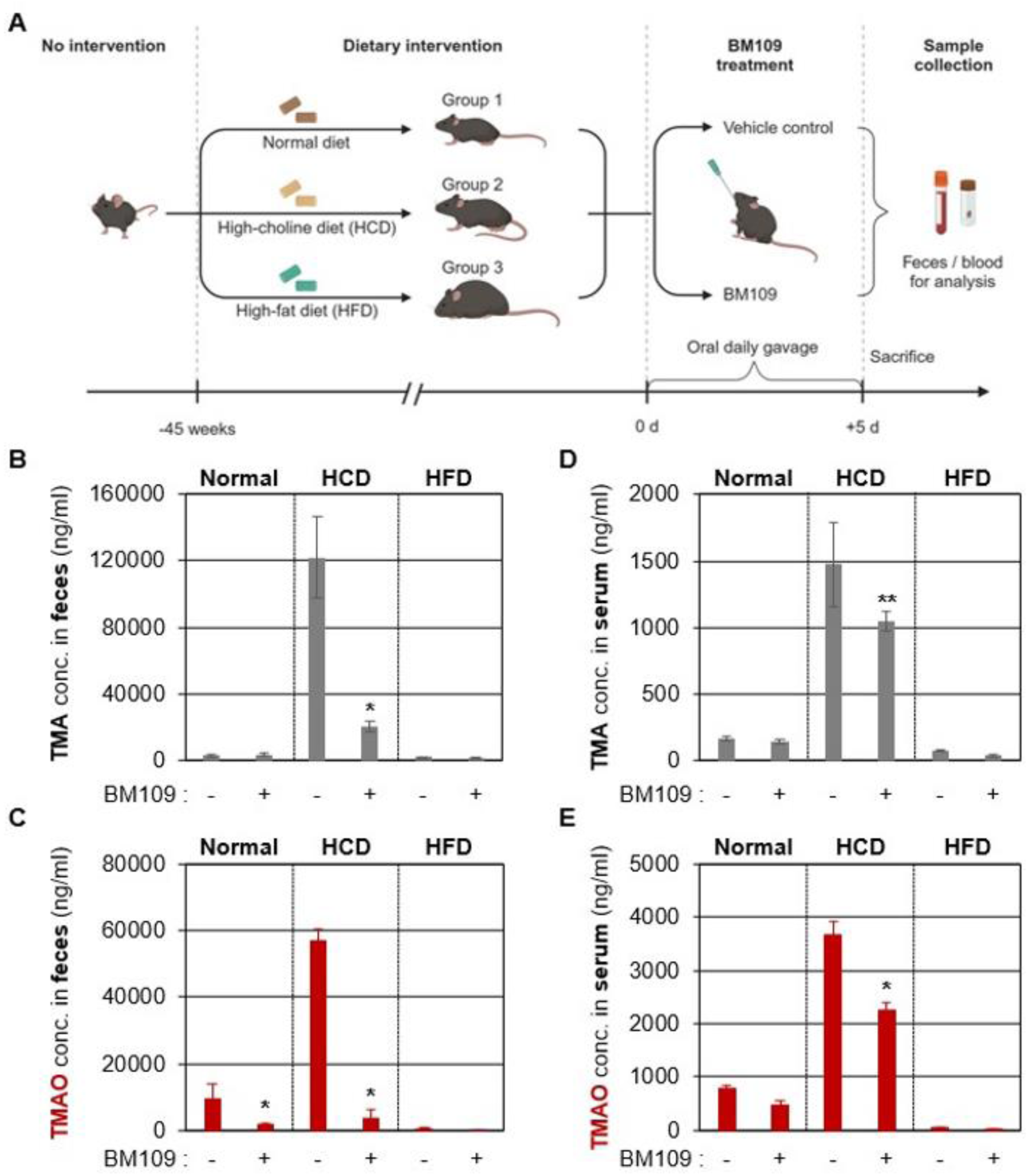
Effects of oral treatment of BM109 cells on changes in TMA and TMAO levels in mouse models. (**A**) Mice were assigned to three dietary groups: normal diet (ND), high-choline diet (HCD), and high-fat diet (HFD), which they were fed for 45 weeks. Each group was subsequently divided into two subgroups: one receiving a vehicle control and the other BM109 (5×10^8^ CFU/day) for 5 consecutive days via oral gavage. (**B, C**) TMA and TMAO concentrations in fecal samples. (**D, E**) TMA and TMAO concentrations in serum samples. Data are expressed as mean ± SD (n=5 per group). In each bar, values of mean ± SD are presented. \**p* < 0.001 vs. control treatment, \*\**p* < 0.05 vs. control treatment.

As shown in Fig. 4B–E, HFD had no significant effect on TMA or TMAO levels. In contrast, HCD feeding markedly increased both metabolites in feces and serum, establishing a suitable model to assess the effects of BM109. In HCD-fed mice, fecal TMA levels were significantly elevated and were reduced by 83% following five days of BM109 administration (Fig. 4B). Similarly, fecal TMAO levels were substantially decreased after BM109 treatment (Fig. 4C), indicating efficient degradation of TMA within the intestinal environment.

We next examined TMA and TMAO levels in mouse serum to assess systemic effects of BM109. In HCD-fed mice, the TMAO concentration reached ∼3,666 ng/mL (Fig. 4E), while the TMA level was ∼1,473 ng/mL (Fig. 4D). Interestingly, the ratio of TMAO to TMA in serum was opposite to that observed in feces (∼56,931 ng/mL vs. ∼121,724 ng/mL). This result further confirms that a significant amount of blood TMAO is produced from the hepatic enzyme flavin-containing monooxygenase 3 (FMO3) using intestinal TMA absorbed through the portal vein and delivered to the liver (*25*). Upon daily treatment with BM109 cells, blood levels of TMA and TMAO decreased by ∼30% and ∼38%, respectively, in the HCD-fed group (Fig. 4D and E). Furthermore, TMAO levels in the Normal diet group also decreased by∼40% after BM109 treatment (Fig. 4E).

Notably, BM109 retained its TMA-degrading activity *in vivo* without supplementation of exogenous electron acceptors, suggesting that the gut environment provides sufficient conditions, potentially including endogenous or diet-derived nitrate to support its metabolic function. Collectively, these findings demonstrate that BM109 effectively degrades intestinal TMA and reduces circulating TMAO levels, supporting its therapeutic potential for TMAO-associated metabolic disorders.

### Neuroprotective Effects of BM109 in the tMCAO Rat Model

Building upon the findings in the HCD diet mouse model, we next evaluated the therapeutic efficacy of BM109 in a disease-specific context using a transient middle cerebral artery occlusion (tMCAO) rat model. This experiment aimed to demonstrate BM109’s potential in mitigating ischemic injury associated with elevated TMAO levels. As illustrated in Fig. 5A, rats were divided into groups based on diet and treatment conditions. BM109 was administered orally for seven days prior to the tMCAO surgery.

**Figure 5.**
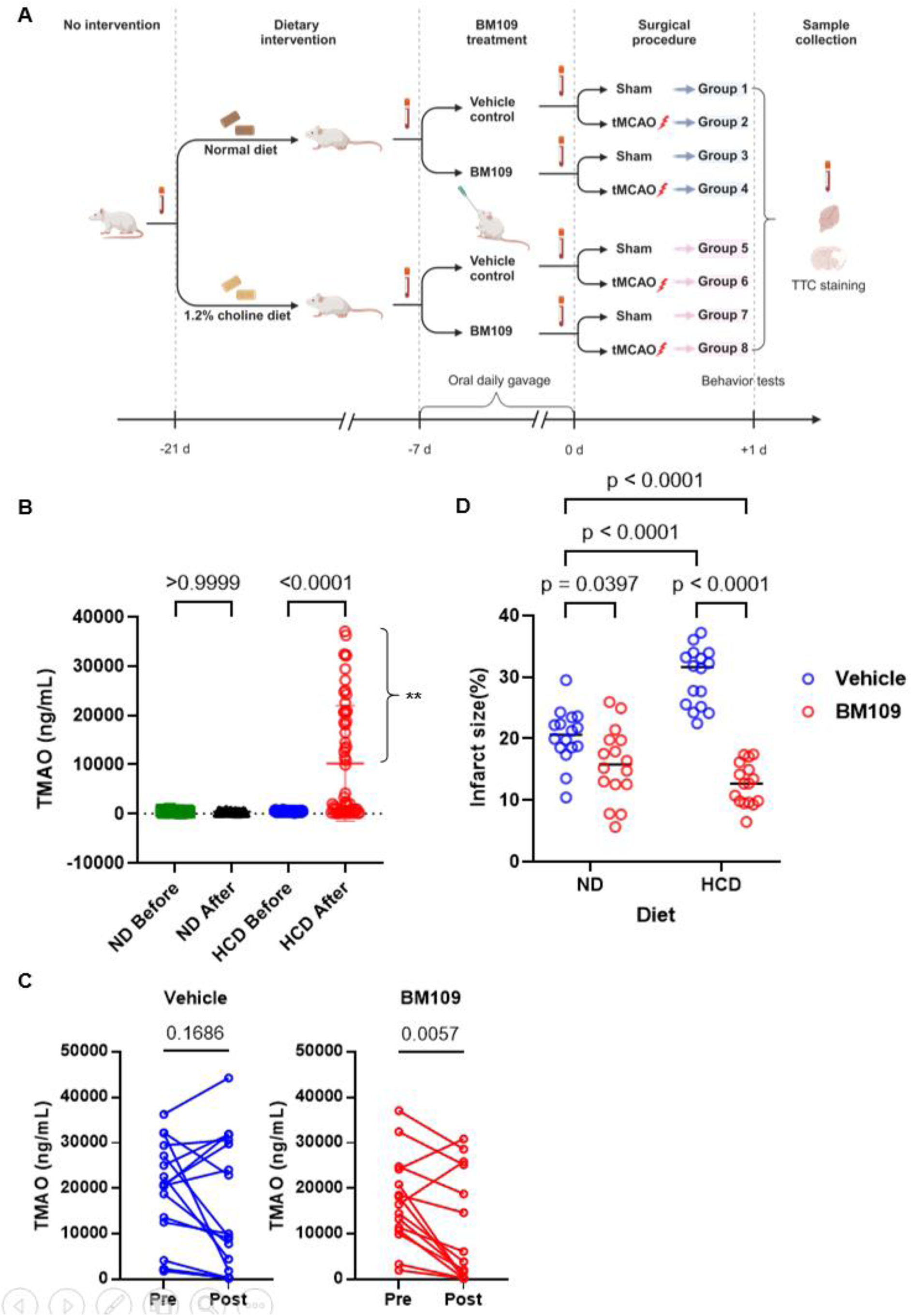
Neuroprotective effects of BM109 in a transient middle cerebral artery occlusion (tMCAO) model. (**A**) Experimental timeline and group assignments. Male Wistar rats (8 weeks old) were fed either a normal diet (ND) or high-choline diet (HCD) for 3 weeks, with BM109 (5×10¹⁰ CFU/day) or vehicle control (15% glycerol) administered during the final 7 days. On the day of surgery, animals underwent a 90-minute tMCAO procedure and were sacrificed 24 h post-reperfusion for infarct analysis and behavioral assessments. (**B**) Plasma TMAO levels at baseline and after dietary intervention. TMAO levels were analyzed in the normal chow (n=60) and HCD (n=58) groups. Statistical analysis was performed using one-way ANOVA. Among the HCD group, 33 rats exhibited more than a threefold increase in TMAO levels compared to baseline (** in the figure). (**C**) Effect of BM109 on plasma TMAO levels in high-TMAO rats (n=33) from panel B (** in the figure). These rats were divided into a vehicle group (n=18) and a BM109-treated group (n=15). Pre- and post-gavage TMAO levels are shown for each group. BM109 significantly reduced TMAO levels compared to the vehicle group, with p-values indicated (paired t-test). (**D**) Infarct size (%) measured by triphenyltetrazolium chloride (TTC) staining to evaluate ischemic brain injury. Control and BM109-treated groups are shown in blue and red, respectively. Statistical significance was determined using paired t-test, with p-values indicated above the comparisons.

To assess the impact of diet on blood TMAO levels, we measured TMAO concentrations after two weeks of feeding either a normal diet or HCD. HCD-fed animals exhibited a marked increase in blood TMAO levels compared to their baseline levels before HCD feeding (Fig. 5B). This trend was consistent with observations in mice (Fig. 4E).

Notably, among the 58 animals fed an HCD, 33 exhibited a substantial increase in blood TMAO levels, with at least a three-fold elevation, as indicated by the double asterisks in Fig. 5B. To further evaluate whether BM109 treatment could lower systemic TMAO levels, we analyzed this subset of animals with markedly elevated TMAO. As depicted in Fig. 5C, BM109 administration resulted in a statistically significant reduction in blood TMAO levels (p = 0.0057), whereas the control group showed no significant changes (p = 0.1686). These findings further support the *in vivo* efficacy of intestinally delivered BM109 in reducing diet-induced TMAO accumulation.

Given this effect, we next examined how this reduction influenced the outcomes of tMCAO. As shown in Fig. 5D, BM109 treatment led to a significant reduction in infarct size across both diet groups. In animals fed a normal diet, BM109 administration reduced infarct size by approximately 22% compared to controls (p = 0.049). This effect was even more pronounced in HCD-fed rats, where BM109 treatment decreased infarct size by ∼58% (30.01 ± 4.67 vs. 12.59 ± 3.40, p < 0.0001) compared to PBS-treated controls. These results highlight the potential of BM109 as a therapeutic strategy for mitigating TMAO-induced ischemic injury. Representative TTC-stained images from each group further illustrate these differences, visually highlighting the neuroprotective effects of BM109 (Fig. S2).

Behavioral assessments further validated BM109’s efficacy. Garcia’s scores revealed significant motor and sensory improvements in BM109-treated rats, particularly in the HCD group (p = 0.0003, Fig. 6A). While Longa’s scores did not show significant differences (Fig. 6B), the modified neurological severity score (mNSS) confirmed BM109’s role in reducing neurological deficits. Specifically, mNSS scores were significantly lower in BM109-treated HCD-fed rats compared to PBS-treated controls (p = 0.0025, Fig. 6C). Together, these findings demonstrate that BM109 effectively reduced HCD-induced TMAO elevation, leading to substantial neuroprotection in the tMCAO model. By lowering systemic TMAO and mitigating ischemic injury, BM109 shows promise as a therapeutic agent for TMAO-associated ischemic stroke.

**Figure 6.**
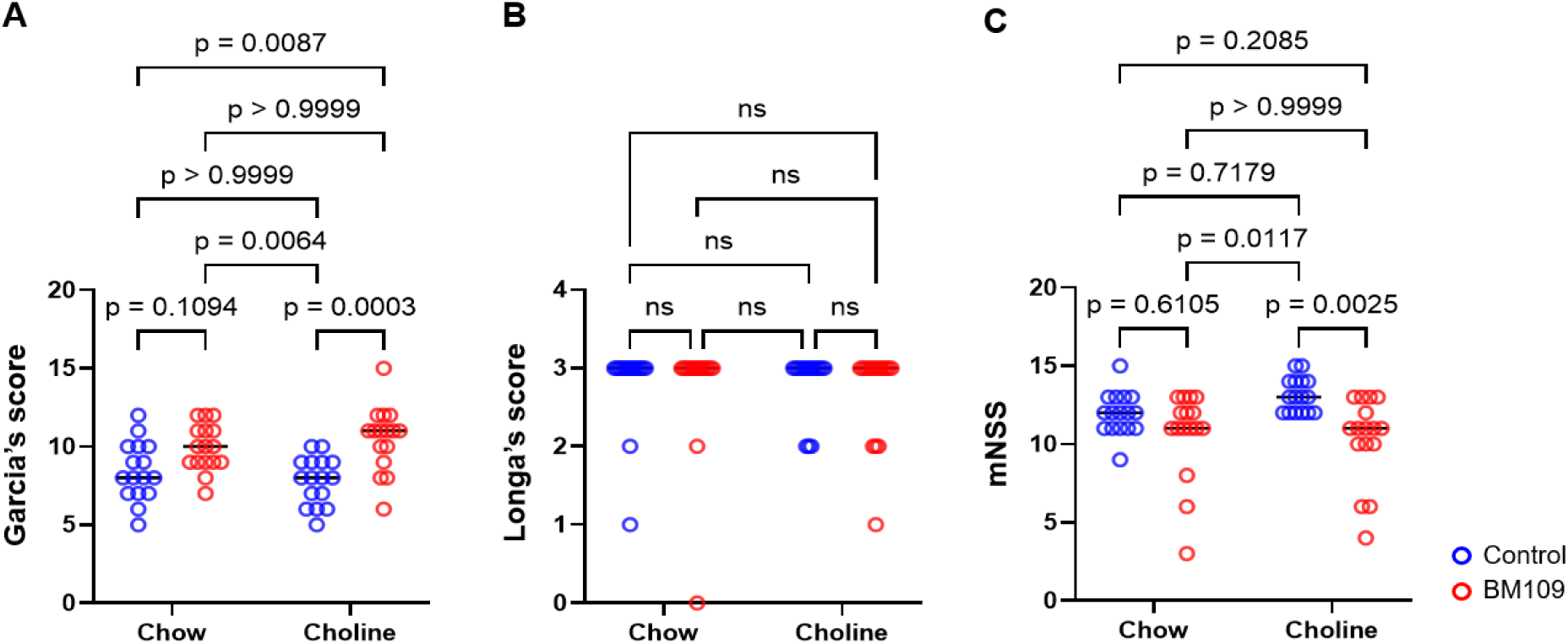
BM109 improves neurological outcomes in a (tMCAO) model. (**A**) Garcia score assessed sensorimotor function and neurological performance. (**B**) Longa score evaluated stroke severity based on motor deficits. (**C**) Modified Neurological Severity Score (mNSS) assessed overall neurological impairment, including motor, sensory, and reflexive functions. In all panels, red circles represent the control group, and blue circles represent the BM109-treated group. Black horizontal lines indicate the mean values for each group. p-values are directly indicated in the figure for statistical comparisons.

### Biosafety assessment of BM109

To evaluate BM109 as a potential live biotherapeutic product (LBP), comprehensive biosafety assessments were conducted. First, hemolytic activity was examined, and BM109 showed no hemolysis on blood agar (Fig. S3A). Second, gastrointestinal safety was assessed in mice administered either PBS or BM109 (n=5). No signs of inflammation, fluid accumulation, or tissue damage were observed, and colonic length remained unchanged (Fig. S3B). Histological analysis further confirmed BM109’s safety, as H&E staining revealed no significant differences in epithelial integrity, crypt architecture, or inflammatory cell infiltration between PBS- and BM109-treated groups (Fig. S3C). Finally, antibiotic resistance was evaluated against eight clinically and agriculturally relevant antibiotics, including ampicillin, streptomycin, and vancomycin (Fig. S4). BM109 was sensitive to all tested antibiotics, although slight resistance to vancomycin was observed. However, considering that BM109 is a Gram-negative bacterium, this is not a significant concern, as vancomycin primarily targets Gram-positive organisms (*47*). Collectively, these results confirm BM109’s safety as an oral LBP, demonstrating no concerning antibiotic resistance and strong potential for mitigating TMA- and TMAO-associated disorders.

## Discussion

TMAO has been suggested as an independent risk factor and effective prognostic marker for various cardiovascular and cerebrovascular diseases. In prospective observational studies conducted in many countries, patients with higher blood TMAO levels invariably exhibited a higher tendency to develop adverse long-term cardiovascular risk in the USA (*29, 48*), Korea (*30*), Japan (*34*), China (*49*), Norway (*50*) and Spain (*51*). Notably, TMAO showed a stronger correlation power than other blood test items, such as LDL-cholesterol and triglycerides, predicting future CVD events (*29, 30, 48, 49*). While the predictive power of TMAO differs based on race of the target patients (*48, 52*), these results demonstrate that TMAO is a metabolite whose amounts need to be decreased for the prevention and treatment of CVD.

TMA, a precursor of TMAO, is enzymatically produced by commensal bacteria that reside in the intestine. Choline-TMA lyase and carnitine oxygenase encoded by *cutC* and *cntA* genes are mainly involved in this process (*53, 54*). Based on comprehensive bioinformatics analysis, 1,107 and 6,783 out of 67,134 bacterial genomes contain *cutC* and *cntA* genes, respectively (*37*). This analysis indicated that the *cutC* gene, in particular, has a very low distribution across the entire bacterial kingdom. However, quantitative gene amplification assays using 50 different human stool samples revealed that the *cutC* gene was detected in all samples, whereas the *cntA* gene was detected in 26% of the samples (*37*). This result suggests that the human intestine requires the presence of *cutC* and *cntA* genes of microbial origin, with *cutC* at a higher frequency, to facilitate enhanced digestion during the process of establishing symbiosis. From an ecological and evolutionary perspective, this observation suggests that the human microbiome has co-evolved toward a symbiotic relationship that favors TMA production, reflecting its adaptive significance in optimizing digestive processes and nutrient metabolism.

Efforts to inhibit microbial TMA production have garnered significant attention due to its contribution to systemic TMAO formation. Among these, a structural analog of choline, 3,3-dimethyl-1-butanol (DMB), has been extensively studied for its suppressive activity against choline-TMA lyase (*55*). In animal studies, DMB treatment led to decreased TMAO levels and subsequent amelioration of TMAO-induced adverse phenotypes (*56, 57*). However, repeated treatment with DMB induces neurotoxic effects in adult mice (*58*). Furthermore, DMB treatment failed to suppress TMA production in an *in vitro* model of human colon fermentation (*59*). More recently, iodomethylcholine (IMC), a mechanism-based inhibitor of choline-TMA lyase, has been shown to effectively reduce TMA and TMAO levels in preclinical models by irreversibly inhibiting microbial TMA formation (*60–62*). However, such approaches rely on broad enzymatic inhibition and may be limited by off-target effects and challenges in selectively modulating complex gut microbial communities. Collectively, these findings suggest that inhibition-based strategies may not fully address TMA production in the complex gut environment. In contrast, our strategy focuses on the direct degradation of TMA by a commensal bacterium, offering a distinct and potentially more targeted approach to reducing systemic TMAO levels.

Lawrence et al. reported an association between circulating bacterial DNA profiles and cardiovascular mortality in a cohort of 405 individuals (*45*). Blood samples obtained from patients who died from CVD contained significantly lower levels of bacterial DNA originating from the genus Paracoccus (p < 0.001). Since circulating bacterial DNA may originate, at least in part, from the translocation of bacteria from the gut, this finding suggests that these patients may have had lower intestinal abundance of Paracoccus. These observations are consistent with our finding that a Paracoccus strain (BM109) exerts beneficial effects on TMA metabolism and cardiovascular-related outcomes. While causality cannot be established, this association raises the possibility that Paracoccus abundance may be linked to cardiovascular health.

Our results (Fig. 4) demonstrate that both TMA and TMAO levels were markedly increased in fecal samples from HCD-fed mice. This observation raises the possibility that TMAO may be present within the intestinal environment, although its precise origin remains unclear. While TMAO is primarily generated in the liver via flavin-containing monooxygenase 3 (FMO3), previous studies have suggested that certain gut bacteria possess TMA monooxygenase activity, which could contribute to local TMAO formation (*63*). Notably, BM109 uniquely encodes TMAO reductase, which catalyzes the reduction of TMAO to TMA, followed by further metabolism via TMA dehydrogenase (Fig. 3). This metabolic capability suggests that BM109 may reduce intestinal TMAO levels, thereby limiting its absorption into the bloodstream.

*Paracoccus* species, including the BM109 strain explored in the current study, are methylotrophic (*64, 65*). Consistent with this metabolic capability, BM109 degraded 10 mM TMA within 24 h *in vitro*, a concentration exceeding typical physiological levels. This high degradation capacity may be supported by downstream metabolic pathways that convert toxic intermediates such as formaldehyde into formate and CO_2_. BM109 exhibited optimal growth at 37 °C and retained activity under anaerobic conditions, supporting its potential functionality within the intestinal environment. Genomic analysis further indicated the presence of genes associated with nitrate-dependent anaerobic respiration, suggesting that BM109 may utilize alternative electron acceptors available in the gut. Together with its complete set of TMA and TMAO metabolic genes, these features support the ability of BM109 to efficiently degrade TMA and TMAO under physiologically relevant conditions.

To evaluate the *in vivo* efficacy of BM109, we employed a high-choline diet (HCD) model, which is known to elevate TMA and TMAO levels. A high-fat diet (HFD) model was also included to assess the impact of dietary fat alone; however, HFD did not significantly alter TMA or TMAO levels. BM109 administration effectively reduced systemic TMAO levels by directly degrading intestinal TMA, thereby intercepting the microbial-host metabolic axis linking gut microbiota to cardiovascular risk. Notably, plasma TMAO levels were consistently decreased under both HCD and normal diet conditions following oral administration of BM109. Short-term treatment (5∼7 days) was sufficient to achieve significant reductions in TMAO levels and was associated with improved outcomes in the tMCAO model. These findings support the potential of BM109 as a microbiome-based therapeutic strategy for reducing systemic TMAO levels. Future studies are warranted to evaluate the long-term efficacy and durability of BM109 treatment *in vivo*.

Previous studies have reported a positive correlation between circulating TMAO levels and infarct severity in experimental stroke models (*66*). Consistent with these findings, our rat experiments demonstrated a relationship between elevated TMAO levels and increased infarct size, which was most pronounced under high-choline diet (HCD) conditions. In these models, dietary choline intake increased plasma TMAO levels, recapitulating TMAO-associated ischemic risk observed in humans. Prior work has also shown that enhancing microbial TMA production, for example through microbiota transplantation enriched in choline-utilizing pathways, exacerbates ischemic injury, whereas disruption of TMA-producing enzymes reduces TMAO levels and improves outcomes. While these approaches primarily aim to reduce TMAO levels by limiting microbial TMA production, our strategy is fundamentally distinct. BM109 directly degrades TMA after its formation, thereby reducing the substrate available for hepatic conversion to TMAO. This mechanism targets the upstream microbial metabolite pool, offering a complementary and potentially more efficient approach to lowering systemic TMAO levels. By enzymatically eliminating TMA, BM109 represents a microbiome-based strategy to mitigate TMAO-associated ischemic injury.

Integration of these findings with prior studies supports the potential of BM109 as a microbiome-based therapeutic strategy. In addition to lowering circulating TMAO levels, BM109 conferred significant neuroprotective effects in ischemic models, linking metabolic modulation to improved disease outcomes. Given the high risk of stroke recurrence (*67, 68*), targeting TMAO may represent a viable approach for reducing recurrent ischemic events. Accordingly, BM109 may have therapeutic potential not only in acute settings but also in long-term risk management. Further studies are warranted to evaluate its durability and clinical applicability in both primary and secondary prevention of stroke.

Although BM109 was isolated from human feces, members of the genus Paracoccus are commonly associated with environmental niches such as soil (*69*). The use of non-traditional or environmental bacteria for therapeutic purposes has been increasingly explored, particularly when a defined mechanism of action can be demonstrated (*70*). In such contexts, functional activity, together with safety considerations, is a key determinant of therapeutic potential. Our findings demonstrate that BM109 effectively degrades TMA and reduces circulating TMAO levels *in vivo*, supporting its potential as a microbiome-based strategy for modulating TMAO-associated cardiovascular and cerebrovascular risk. Further studies are warranted to evaluate its long-term efficacy and clinical applicability.

## Methods

### Ethical Statements

All animal experiments in this study were performed in compliance with the guidelines established by the Department of Animal Resources at the Yonsei Biomedical Research Institute. This study was approved by the Institutional Animal Care and Use Committee (IACUC) of Yonsei University College of Medicine under permit numbers 2024-0143 and 2021-0139 and 2023-0063.

### Screening human fecal microbiome for bacteria with TMA-degrading capabilities

A collection of human feces (Severance Hospital, IRB approval number, 4-2016-0850) was used to yield pools of microbiome suspensions. Aliquots of fecal bacterial suspensions were inoculated onto agar plates of DLB (Diluted Luria Bertani) plus 7% MeOH. DLB medium was prepared by 20% dilution of LB medium [1% (w/v) tryptone, 0.5% (w/v) yeast extract, and 1% (w/v) sodium chloride] in water. At 48 h post-inoculation, the colonies were recovered as potential strains with methylotrophic features. Each colony was then inoculated in a broth medium containing DLB+5 mM TMA to examine its growth and TMA-degrading capability. After cultivation, the spent medium was collected for TMA quantification. Potential TMA degraders were identified by sequencing the entire region of the 16S rRNA gene.

### TMA degradation challenge with additional nutrition

To determine whether the BM109 strain could still degrade TMA in the presence of additional carbon and nitrogen sources, BM109 was cultivated in LB broth supplemented with 10 mM TMA. NaNO_3_ (60 mM) was added as an alternative electron acceptor to stimulate anaerobic growth. For aerobic incubation, BM109 cells were grown by shaking (180 rpm) at 37 °C. Anaerobic cultivation was achieved in an anaerobic chamber (Coy Lab Products, Grass Lake, MI), where a gas mix (5% H_2_ and 95% N_2_) was used to fill the atmosphere. BM109 was grown statically inside an anaerobic chamber. We harvested the aerobic culture after 24 h and the anaerobic culture after 48 h to quantify the residual TMA. We also cultured *Paracoccus denitrificans* ATCC17741 (PD) using the same protocol as a negative control. OD_600nm_ values of each bacterial culture were measured after incubation.

### Colorimetric TMA quantification assay

Colorimetric TMA quantification was performed using a modified picric acid-toluene assay (*71, 72*). Serially diluted TMA standards (10∼1.25 mM) were prepared in LB for the standard curve. BM109 and PD cultures were centrifuged, and supernatants were processed with formaldehyde, toluene, and potassium carbonate. After vortexing and shaking, the toluene phase was mixed with picric acid-toluene solution, and absorbance at 410 nm was measured using a microplate reader.

### Whole-genome sequencing of BM109 and genome analysis

Genomic DNA was purified using G-spin Total DNA Extraction kit (iNtRON). Genome sequencing was performed using PacBio Sequel and Illumina NovaSeq6000 platform. DNA is required to prepare size-selected approximately 15 kb SMRTbell templates with barcoded overhang adapters. For PacBio Sequel sequencing, library was prepared using PacBio SMRTbell Express Template Prep kit 2.0. The Sequel Sequencing Kit 3.0 and SMRT cells 1M v3 Tray was used for sequencing. SMRT cells (Pacific Biosciences) using 600 min movies were captured for each SMRT cell using the PacBio Sequel (Pacific Biosciences) sequencing platform by Macrogen (Seoul, Korea). Reads from PacBio Sequel system were assembled using SMRTlink 10.1.0.119588 (*73*). Illumina raw reads were used for error correction. The assembly was corrected using high quality Illumina reads by Pilon v1.21 (*74*). The whole-genome sequence data for BM109 have been deposited in GenBank under accession number PRJNA891425.

To investigate the TMAO metabolic pathway in the BM109 genome and its closely related species, we employed a comprehensive analytical approach. Gene prediction and annotation were performed using Prokka (version 1.14), InterProScan (version 5.30-69), and PSI-BLAST (version 2.4.0), with the EggNog database (version 4.5) serving as a reference for functional classification. To examine the BM109 genome in comparison with closely related species, we retrieved genome sequences of related species from the NCBI database. These genomes were subsequently analyzed using BlastKOALA, which enabled the identification of genes relevant to the TMAO pathway.

The identified genes from these genomes were then aligned with the gene set of BM109, allowing for the identification of conserved elements across species. This comparative analysis provided key insights into the genetic basis of the TMAO pathway in BM109 and its evolutionary relatives, highlighting conserved genetic features that may underlie the functionality and adaptability of this pathway.

### Chronic Dietary Model in Mice

For the chronic dietary model, C57BL/6J female mice (5 weeks old) were divided into three groups of ten mice each and fed one of the following diets for 45 weeks: a chemically defined normal-chow diet (NC, 0.08% wt/wt total choline), the same chemically defined diet supplemented with 1% wt/wt choline (HCD, Envigo TD.130328), or a chemically defined 60% wt/wt fat diet (HFD, Envigo TD.06414). Female mice were used in this study because hepatic expression of flavin-containing monooxygenase 3 (FMO3), the key enzyme responsible for TMAO production, is markedly reduced in male mice, resulting in significantly lower TMAO levels (*75*). After 45 weeks, each group was split into subgroups of 5 mice. Control groups were gavaged with 100 μL of tap water daily for five consecutive days. Experimental groups received 100 μL of BM109 suspension, which was prepared by harvesting activated BM109 cells after 24 h of incubation, concentrating them to 5×10⁹ CFU/mL, washing the pellet, and resuspending it in PBS before administration.

### Transient Middle Cerebral Artery Occlusion (tMCAO) Model in Rats

For the tMCAO model, male Wistar rats (8 weeks old) were randomly assigned to eight groups based on diet (chow or choline-supplemented) and treatment (BM109 or vehicle) with or without tMCAO surgery. The study followed ARRIVE guidelines, and sample size was determined using MedCalc software. Focal ischemia was induced via tMCAO under isoflurane anesthesia following procedures described previously (*76*). A monofilament was introduced through the external carotid artery to occlude the middle cerebral artery for 90 minutes, after which reperfusion was allowed. Sham-operated controls underwent the same procedure without occlusion. Neurological function was assessed 24 h post-surgery using the Garcia test (*77*), Longa score (*78*), and modified Neurological Severity Score (mNSS) (*79*). Brains were collected 24 h post-surgery, sectioned, and stained with TTC. Infarcted areas were quantified using image analysis software, calculating infarct volume as a percentage of total brain area (*80*). Samples were collected at four time points: before diet administration, before BM109 supplementation, before surgery, and before euthanasia. Plasma was isolated for TMAO quantification.

### Quantification of TMA and TMAO

Blood and fecal samples in Figure 4 were collected 24 h after the final gavage. Blood plasma was obtained by centrifugation, while fecal samples were homogenized in PBS, centrifuged, and the supernatant was collected. Both plasma and fecal supernatants were diluted in methanol (1:4 vol/vol), centrifuged, and the supernatants were analyzed for TMA and TMAO using liquid chromatography-mass spectrometry (LC-MS) at the National Instrumentation Center for Environmental Management (NICEM), Seoul National University. The results were compared across six groups (NC, HC, HF), each with control and experimental subgroups. Additionally, TMAO quantification in rat blood samples was conducted at the Department of Laboratory Medicine, Severance Hospital following established procedures (*30*).

### Assessment of BM109 safety

The hemolytic activity of BM109 was assessed by streaking the strain on a blood agar plate, as described in a previous study (*81*). Intestinal and colon tissues were collected from C57BL/6J mice following oral administration of BM109 (5×10⁸ CFU/day for five days) or PBS. The tissues were fixed in 4% paraformaldehyde, embedded in paraffin, and sectioned for histological analysis. Hematoxylin and eosin (H&E) staining was performed to evaluate potential tissue damage, inflammation, or histological abnormalities, following the methods outlined elsewhere (*82*). The antibiotic susceptibility of BM109 was evaluated using the E-test method, following the Clinical and Laboratory Standards Institute (CLSI) guidelines (*83*).

### Statistical analysis

Statistical analyses were performed using GraphPad Prism (version 10.4.1, GraphPad Software, San Diego, CA, USA). Data are presented as mean ± standard deviation (SD), and a p-value < 0.05 was considered statistically significant. Differences between paired samples were analyzed using a paired t-test, while comparisons between multiple groups were assessed using one-way and two-way ANOVA with Bonferroni post hoc tests. Repeated measures mixed models were applied where appropriate.

## Supporting information

Genome sequence information

## Acknowledgements

This work was supported by grants from the National Research Foundation (NRF) of Korea, funded by the Korean Government (2022M3A9F3017506 and 2019R1A6A1A03032869). This research was supported by Korea Drug Development Fund funded by Ministry of Science and ICT, Ministry of Trade, Industry, and Energy, and Ministry of Health and Welfare (RS-2024-00443597). This research was also supported by the National Institute of Health (NIH) research project (2024-ER2107-00). This work was supported by the National Research Foundation of Korea (NRF) grant funded by the Korea government (MSIT) (No. 2022R1A2C1007948).

## Author contributions

JSY, CEY, and JBK contributed equally as co-first authors. All authors participated in experimental design and data interpretation. JSY, CEY, JBK, MAA, and SGK performed the experiments and analyses. CEY, YBK, and HSN conducted the *in vivo* stroke model experiments and behavioral assessments. JSY, CEY, JBK, and SSY drafted the manuscript. HSN and SSY supervised the study as co-corresponding authors.

---

**Figure S1.**
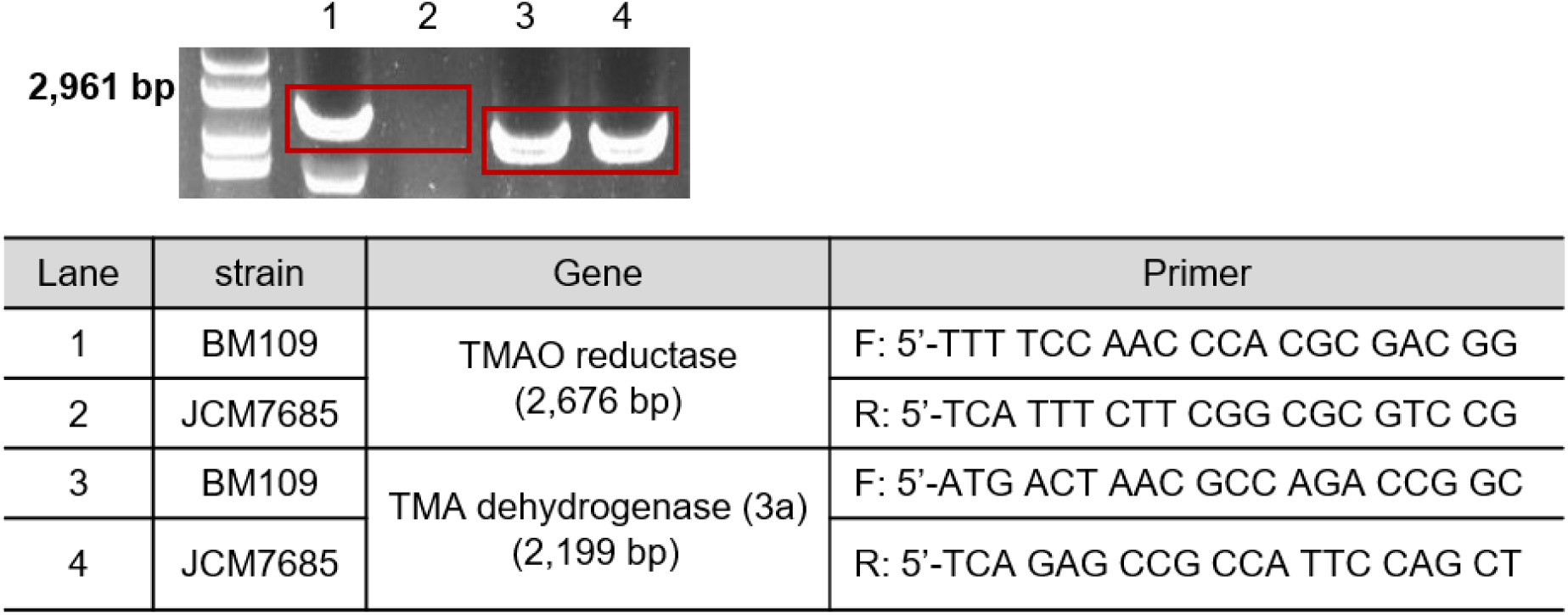
PCR confirmation of the gene encoding TMAO reductase in BM109, but not in JCM7685. Genomic DNA extracted from BM109 (lanes 1 and 3) or JCM7685 (lanes 2 and 4) was PCR amplified for the presence of the TMAO reductase-encoding gene. Primer sequences used for PCR reactions are shown in the table.

**Figure S2.**
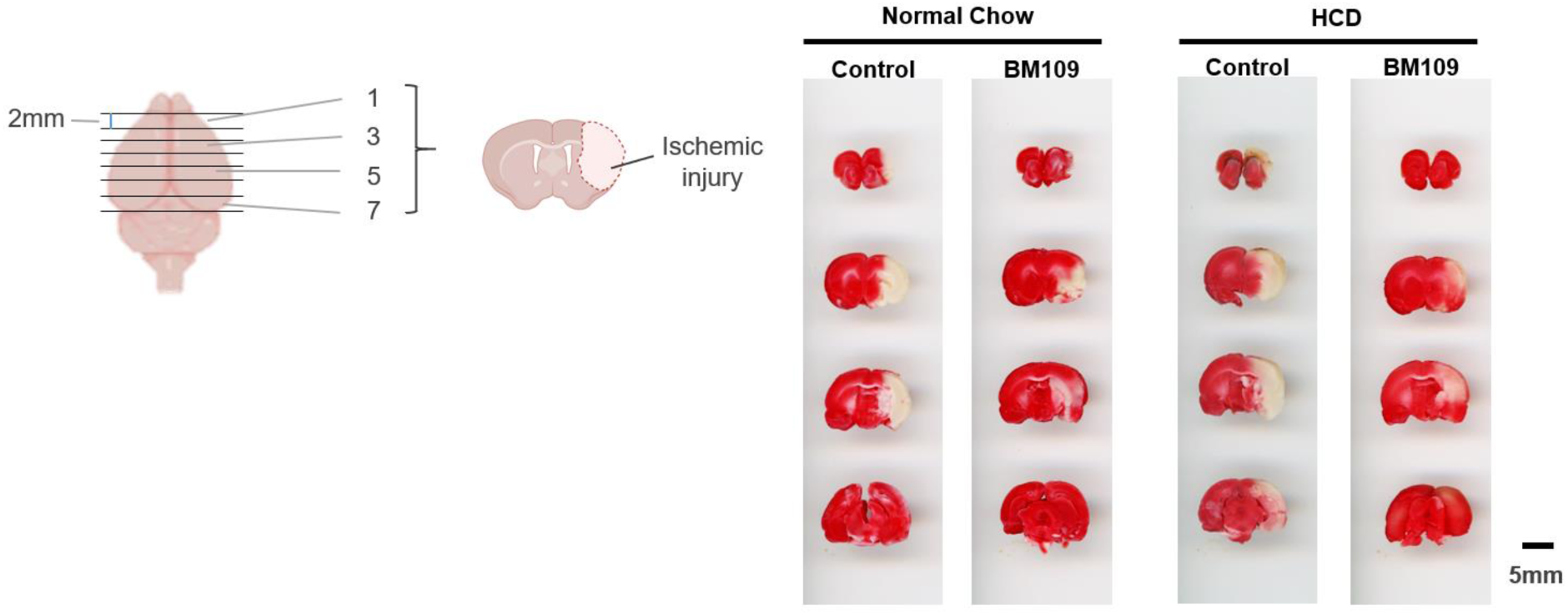
BM109 reduces ischemic injury in a tMCAO model. Representative coronal brain sections stained with 2,3,5-triphenyltetrazolium chloride (TTC) to visualize infarcted areas 24 h post-reperfusion. The schematic on the left illustrates the sectioning method, where four coronal slices (1, 3, 5, and 7) were obtained at 2-mm intervals. White areas indicate infarcted (ischemic) tissue, while red regions represent viable tissue. Brain sections are shown for rats fed a normal chow diet (left) or a high-choline diet (HCD, right), with control (vehicle-treated) and BM109-treated groups. BM109 administration visibly reduced infarct size under both dietary conditions. Scale bar = 5 mm. The schematic illustration on the left was created using BioRender.com.

**Figure S3.**
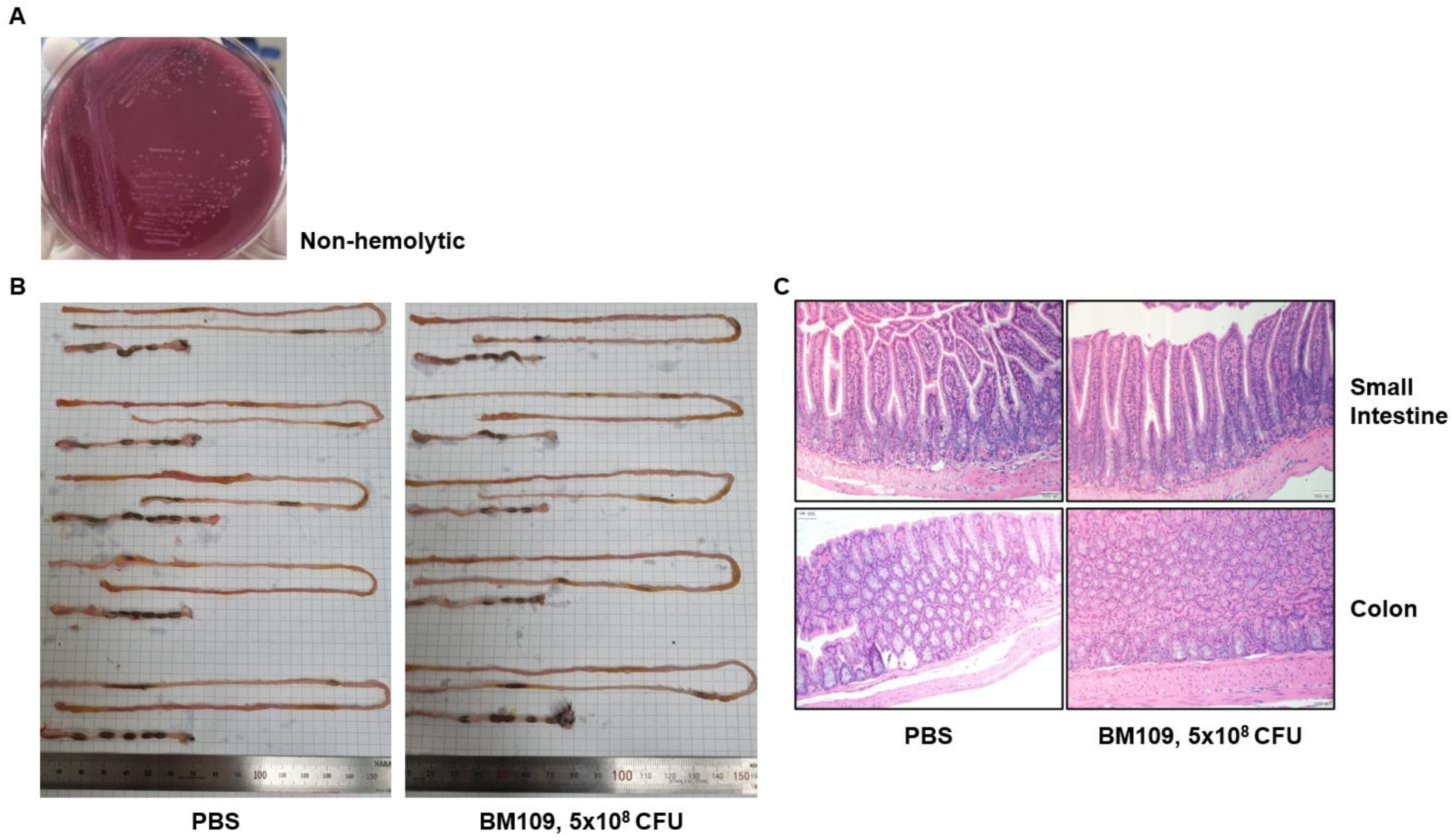
Safety assessment of BM109 administration *in vivo*. **(A)** Hemolysis assay on blood agar plate showing that BM109 is non-hemolytic, indicating no hemolytic activity. **(B)** Gross examination of the intestinal tract from mice administered either PBS (left) or BM109 (5×10⁸ CFU, right). No visible abnormalities, inflammation, or structural damage were observed in the BM109-treated group. **(C)** Histological analysis of the small intestine and colon following BM109 administration. Representative hematoxylin and eosin (H&E)-stained sections of the small intestine (top) and colon (bottom) show no significant histopathological differences between the PBS and BM109-treated groups.

**Figure S4.**
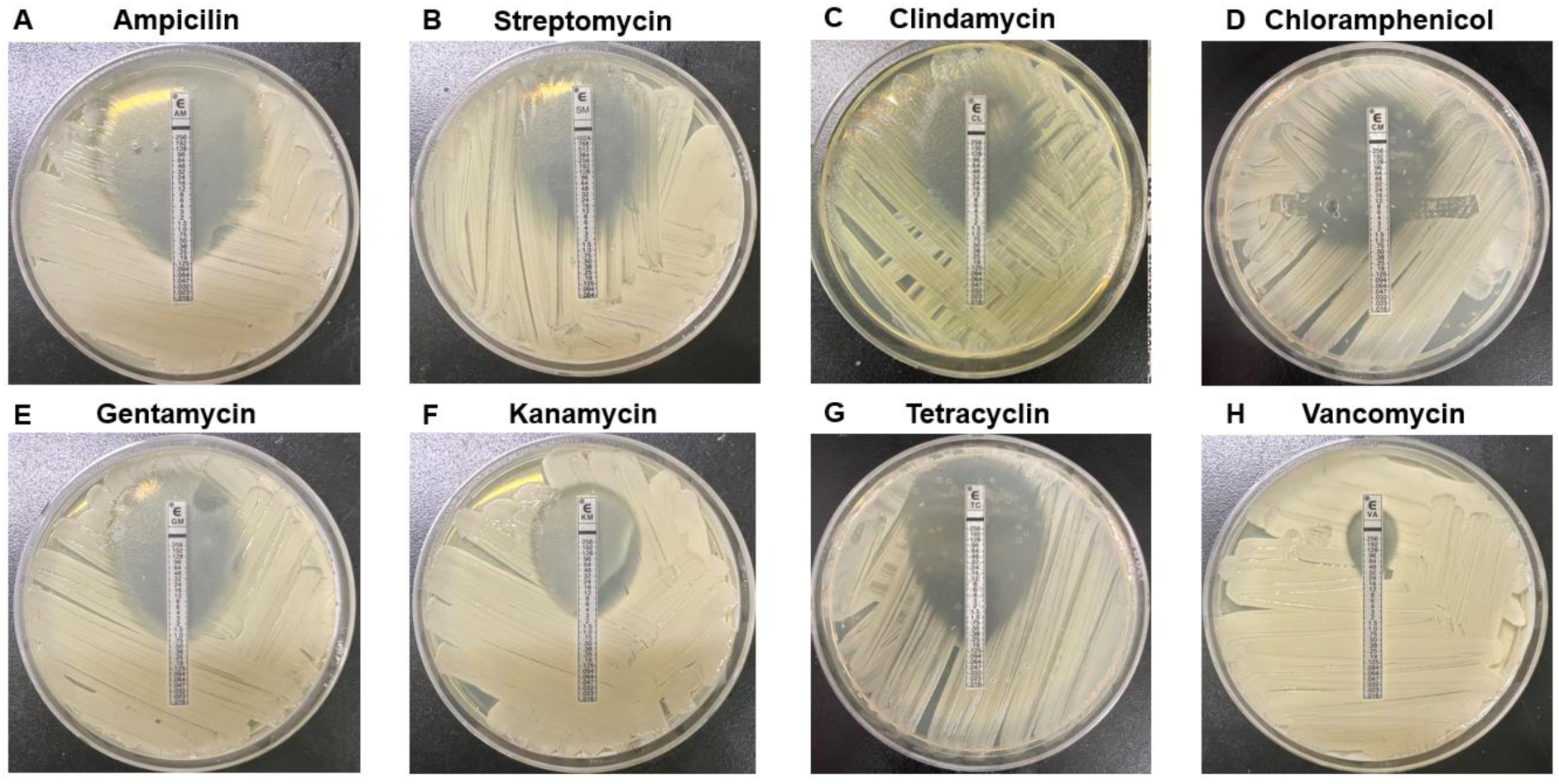
Antibiotic susceptibility testing of BM109. E-test strips were used to determine the minimum inhibitory concentrations (MICs) of BM109 against eight clinically and agriculturally relevant antibiotics. BM109 was streaked onto Müller-Hinton agar plates, and E-test strips impregnated with a gradient of each antibiotic were applied. The inhibition zones indicate BM109’s susceptibility to each antibiotic. The tested antibiotics include ampicillin (AM), streptomycin (SM), chloramphenicol (CL), clindamycin (CM), gentamicin (GM), kanamycin (KM), tetracycline (TC), and vancomycin (VA). BM109 exhibited susceptibility to all tested antibiotics, with slight resistance observed for vancomycin, which is expected for Gram-negative bacteria.

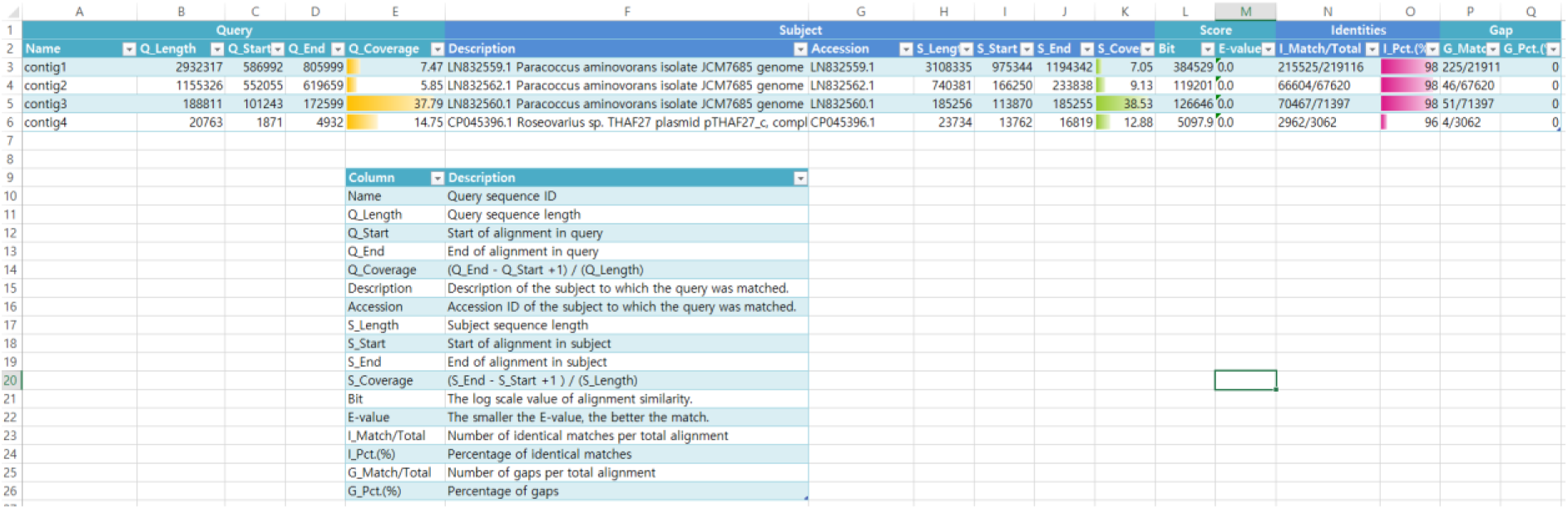

Supplementary information. Genome alignment summary of BM109 contigs with reference *Paracoccus aminovorans* genomes and plasmid sequences

## References

1. J. S. You et al., Commensal-derived metabolites govern Vibrio cholerae pathogenesis in host intestine. Microbiome 7, 132 (2019).

2. M. Y. Yoon et al., A single gene of a commensal microbe affects host susceptibility to enteric infection. Nat Commun 7, 11606 (2016).

3. R. Sender, S. Fuchs, R. Milo, Revised Estimates for the Number of Human and Bacteria Cells in the Body. PLoS Biol 14, e1002533 (2016).

4. E. Bianconi et al., An estimation of the number of cells in the human body. Ann Hum Biol 40, 463–471 (2013).

5. Y. Ding, Z. Zhang, K. Wang, C. Jiang, The microbiome regulates host metabolic health and diseases through microbial enzymes. Nat Rev Gastroenterol Hepatol, (2026).

6. Y. Y. Ma, X. Li, J. T. Yu, Y. J. Wang, Therapeutics for neurodegenerative diseases by targeting the gut microbiome: from bench to bedside. Transl Neurodegener 13, 12 (2024).

7. M. Wang et al., The Gut Microbial Metabolite Trimethylamine N-oxide, Incident CKD, and Kidney Function Decline. J Am Soc Nephrol 35, 749–760 (2024).

8. Y. Makkieh et al., The gut-heart axis: Exploring the role of the gut microbiome in cardiovascular health - A focused systematic review. Am Heart J Plus 61, 100687 (2026).

9. I. Szegedi, D. Bomberak, Z. Eles, L. Loczi, Z. Bagoly, Cardiovascular disease and microbiome: focus on ischemic stroke. Pol Arch Intern Med 135, (2025).

10. S. Feng, Y. Jiang, J. Jiang, H. Bian, R. Zhu, Effects of diet-modulated gut microbiota and microbial metabolites in atherosclerosis. Biomed Pharmacother 198, 119328 (2026).

11. M. P. Khuu et al., The gut microbiota in thrombosis. Nat Rev Cardiol 22, 121–137 (2025).

12. A. Piccioni et al., Gut Microbiota and Environment in Coronary Artery Disease. Int J Environ Res Public Health 18, (2021).

13. H. X. Yang et al., Gut microbiota-derived butyrate prevents aortic dissection via GPR41. Acta Pharmacol Sin 46, 3230–3243 (2025).

14. K. Kasahara et al., Interactions between Roseburia intestinalis and diet modulate atherogenesis in a murine model. Nat Microbiol 3, 1461–1471 (2018).

15. K. T. Kim et al., Antioxidant and Anti-Inflammatory Effect and Probiotic Properties of Lactic Acid Bacteria Isolated from Canine and Feline Feces. Microorganisms 9, (2021).

16. J. Zhou et al., Fecal Microbiota Transplantation in Mice Exerts a Protective Effect Against Doxorubicin-Induced Cardiac Toxicity by Regulating Nrf2-Mediated Cardiac Mitochondrial Fission and Fusion. Antioxid Redox Signal 41, 1–23 (2024).

17. M. Troseid, G. O. Andersen, K. Broch, J. R. Hov, The gut microbiome in coronary artery disease and heart failure: Current knowledge and future directions. EBioMedicine 52, 102649 (2020).

18. S. Fromentin et al., Microbiome and metabolome features of the cardiometabolic disease spectrum. Nat Med 28, 303–314 (2022).

19. W. H. W. Tang, F. Backhed, U. Landmesser, S. L. Hazen, Intestinal Microbiota in Cardiovascular Health and Disease: JACC State-of-the-Art Review. J Am Coll Cardiol 73, 2089–2105 (2019).

20. Y. Talmor-Barkan et al., Metabolomic and microbiome profiling reveals personalized risk factors for coronary artery disease. Nat Med 28, 295–302 (2022).

21. N. Kazemian, M. Mahmoudi, F. Halperin, J. C. Wu, S. Pakpour, Gut microbiota and cardiovascular disease: opportunities and challenges. Microbiome 8, 36 (2020).

22. R. G. Pushpass, S. Alzoufairi, K. G. Jackson, J. A. Lovegrove, Circulating bile acids asa link between the gut microbiota and cardiovascular health: impact of prebiotics, probiotics and polyphenol-rich foods. Nutr Res Rev, 1–20 (2021).

23. Z. Wang et al., Gut flora metabolism of phosphatidylcholine promotes cardiovascular disease. Nature 472, 57–63 (2011).

24. L. Cui, T. Zhao, H. Hu, W. Zhang, X. Hua, Association Study of Gut Flora in Coronary Heart Disease through High-Throughput Sequencing. Biomed Res Int 2017, 3796359 (2017).

25. S. H. Kim, M. Y. Yoon, S. S. Yoon, TMAO and the gut microbiome: implications for the CVD-CKD-IBD axis. Ann Med 57, 2522324 (2025).

26. A. B. Roberts et al., Development of a gut microbe-targeted nonlethal therapeutic to inhibit thrombosis potential. Nat Med 24, 1407–1417 (2018).

27. L. Xia, Z. Wang, X. Chen, TMAO Induces Vascular Endothelial Cells Pyroptosis Through TET2-CYTB-ROS Pathway. J Inflamm Res 18, 8719–8733 (2025).

28. L. Xiu, P. Zhao, X. Gu, B. Yu, Role of Trimethylamine N-Oxide in Assessing Plaque Instability of the Culprit Lesion in Chinese Patients With ST-Elevation Myocardial Infarction: Insights From a 7-Year Long-Term Follow-Up Study. Clin Transl Sci 18, e70357 (2025).

29. W. H. Tang et al., Intestinal microbial metabolism of phosphatidylcholine and cardiovascular risk. N Engl J Med 368, 1575–1584 (2013).

30. H. S. Nam et al., Elevation of the Gut Microbiota Metabolite Trimethylamine N-Oxide Predicts Stroke Outcome. J Stroke 21, 350–352 (2019).

31. X. Yu et al., Trimethylamine N-oxide predicts cardiovascular events in coronary artery disease patients with diabetes mellitus: a prospective cohort study. Front Endocrinol (Lausanne) 15, 1360861 (2024).

32. T. Suzuki, L. M. Heaney, D. J. Jones, L. L. Ng, Trimethylamine N-oxide and Risk Stratification after Acute Myocardial Infarction. Clin Chem 63, 420–428 (2017).

33. T. Suzuki et al., Association with outcomes and response to treatment of trimethylamine N-oxide in heart failure: results from BIOSTAT-CHF. Eur J Heart Fail 21, 877–886 (2019).

34. Y. Kinugasa et al., Trimethylamine N-oxide and outcomes in patients hospitalized with acute heart failure and preserved ejection fraction. ESC Heart Fail 8, 2103–2110 (2021).

35. T. Shafi et al., Trimethylamine N-Oxide and Cardiovascular Events in Hemodialysis Patients. J Am Soc Nephrol 28, 321–331 (2017).

36. P. Andrikopoulos et al., Evidence of a causal and modifiable relationship between kidney function and circulating trimethylamine N-oxide. Nat Commun 14, 5843 (2023).

37. S. Rath, B. Heidrich, D. H. Pieper, M. Vital, Uncovering the trimethylamine-producing bacteria of the human gut microbiota. Microbiome 5, 54 (2017).

38. K. A. Romano, E. I. Vivas, D. Amador-Noguez, F. E. Rey, Intestinal microbiota composition modulates choline bioavailability from diet and accumulation of the proatherogenic metabolite trimethylamine-N-oxide. mBio 6, e02481 (2015).

39. J. Colby, L. J. Zatman, Trimethylamine metabolism in obligate and facultative methylotrophs. Biochem J 132, 101–112 (1973).

40. C. Loechel, A. Basran, J. Basran, N. S. Scrutton, E. A. Hall, Using trimethylamine dehydrogenase in an enzyme linked amperometric electrode. Part 1. Wild-type enzyme redox mediation. Analyst 128, 166–172 (2003).

41. A. F. Hallett, R. Cooper, Respiratory infection in an intensive care unit. S Afr Med J 52, 1095–1098 (1977).

42. H. Gu et al., A case report of Klebsiella aerogenes-caused lumbar spine infection identified by metagenome next-generation sequencing. BMC Infect Dis 22, 616 (2022).

43. T. Urakami, H. Araki, H. Oyanagi, K. Suzuki, K. Komagata, Paracoccus aminophilus sp. nov. and Paracoccus aminovorans sp. nov., which utilize N,N-dimethylformamide. Int J Syst Bacteriol 40, 287–291 (1990).

44. H. S. Han et al., Correlations of the Gastric and Duodenal Microbiota with Histological, Endoscopic, and Symptomatic Gastritis. J Clin Med 8, (2019).

45. G. Lawrence et al., The blood microbiome and its association to cardiovascular disease mortality: case-cohort study. BMC Cardiovasc Disord 22, 344 (2022).

46. T. K. Le, Y. J. Lee, G. H. Han, S. J. Yeom, Methanol Dehydrogenases as a Key Biocatalysts for Synthetic Methylotrophy. Front Bioeng Biotechnol 9, 787791 (2021).

47. A. Zhou et al., Synergistic interactions of vancomycin with different antibiotics against Escherichia coli: trimethoprim and nitrofurantoin display strong synergies with vancomycin against wild-type E. coli. Antimicrob Agents Chemother 59, 276–281 (2015).

48. Y. Lee et al., Longitudinal Plasma Measures of Trimethylamine N-Oxide and Risk of Atherosclerotic Cardiovascular Disease Events in Community-Based Older Adults. J Am Heart Assoc 10, e020646 (2021).

49. Z. Dong et al., The Association between Plasma Levels of Trimethylamine N-Oxide and the Risk of Coronary Heart Disease in Chinese Patients with or without Type 2 Diabetes Mellitus. Dis Markers 2018, 1578320 (2018).

50. M. Troseid et al., Microbiota-dependent metabolite trimethylamine-N-oxide is associated with disease severity and survival of patients with chronic heart failure. J Intern Med 277, 717–726 (2015).

51. C. Roncal et al., Trimethylamine-N-Oxide (TMAO) Predicts Cardiovascular Mortality in Peripheral Artery Disease. Sci Rep 9, 15580 (2019).

52. Y. Yazaki et al., Ethnic differences in association of outcomes with trimethylamine N-oxide in acute heart failure patients. ESC Heart Fail 7, 2373–2378 (2020).

53. Y. Q. Yang, W. H. Deng, R. Z. Liao, Mechanistic Insights into Choline Degradation Catalyzed by the Choline Trimethylamine-Lyase CutC. J Phys Chem B 129, 5438–5448 (2025).

54. W. K. Wu et al., Gut microbes with the gbu genes determine TMAO production from L-carnitine intake and serve as a biomarker for precision nutrition. Gut Microbes 17, 2446374 (2025).

55. Z. Wang et al., Non-lethal Inhibition of Gut Microbial Trimethylamine Production for the Treatment of Atherosclerosis. Cell 163, 1585–1595 (2015).

56. G. Liu et al., Inhibition of Microbiota-dependent Trimethylamine N-Oxide Production Ameliorates High Salt Diet-Induced Sympathetic Excitation and Hypertension in Rats by Attenuating Central Neuroinflammation and Oxidative Stress. Front Pharmacol 13, 856914 (2022).

57. G. Wang et al., 3,3-Dimethyl-1-butanol attenuates cardiac remodeling in pressure-overload-induced heart failure mice. J Nutr Biochem 78, 108341 (2020).

58. J. Mao et al., Repeated 3,3-Dimethyl-1-butanol exposure alters social dominance in adult mice. Neurosci Lett 758, 136006 (2021).

59. P. Day-Walsh et al., The use of an in-vitro batch fermentation (human colon) model for investigating mechanisms of TMA production from choline, L-carnitine and related precursors by the human gut microbiota. Eur J Nutr 60, 3987–3999 (2021).

60. Y. Tang et al., Intestinal metabolite TMAO promotes CKD progression by stimulating macrophage M2 polarization through histone H4 lysine 12 lactylation. Cell Death Differ 33, 314–326 (2026).

61. W. Zhang et al., Inhibition of microbiota-dependent TMAO production attenuates chronic kidney disease in mice. Sci Rep 11, 518 (2021).

62. P. Pathak et al., Small molecule inhibition of gut microbial choline trimethylamine lyase activity alters host cholesterol and bile acid metabolism. Am J Physiol Heart Circ Physiol 318, H1474–H1486 (2020).

63. D. Fennema, I. R. Phillips, E. A. Shephard, Trimethylamine and Trimethylamine N-Oxide, a Flavin-Containing Monooxygenase 3 (FMO3)-Mediated Host-Microbiome Metabolic Axis Implicated in Health and Disease. Drug Metab Dispos 44, 1839–1850 (2016).

64. J. Czarnecki, D. Bartosik, Diversity of Methylotrophy Pathways in the Genus Paracoccus (Alphaproteobacteria). Curr Issues Mol Biol 33, 117–132 (2019).

65. S. C. Baker et al., Molecular genetics of the genus Paracoccus: metabolically versatile bacteria with bioenergetic flexibility. Microbiol Mol Biol Rev 62, 1046–1078 (1998).

66. W. Zhu et al., Gut microbes impact stroke severity via the trimethylamine N-oxide pathway. Cell Host Microbe 29, 1199–1208 e1195 (2021).

67. J. Xue et al., Residual Risk of Trimethylamine-N-Oxide and Choline for Stroke Recurrence in Patients With Intensive Secondary Therapy. J Am Heart Assoc 11, e027265 (2022).

68. Y. Y. Chen, Z. S. Ye, N. G. Xia, Y. Xu, TMAO as a Novel Predictor of Major Adverse Vascular Events and Recurrence in Patients with Large Artery Atherosclerotic Ischemic Stroke. Clin Appl Thromb Hemost 28, 10760296221090503 (2022).

69. R. Kumar, B. Singh, V. K. Gupta, Biodegradation of fipronil by Paracoccus sp. in different types of soil. Bull Environ Contam Toxicol 88, 781–787 (2012).

70. T. Brodmann et al., Safety of Novel Microbes for Human Consumption: Practical Examples of Assessment in the European Union. Front Microbiol 8, 1725 (2017).

71. S. Mohri, M. Kanauchi, Isolation of Lactic Acid Bacteria Eliminating Trimethylamine (TMA) for Application to Fishery Processing. Methods Mol Biol 1887, 109–117 (2019).

72. X. Heng, W. Liu, W. Chu, Identification of choline-degrading bacteria from healthy human feces and used for screening of trimethylamine (TMA)-lyase inhibitors. Microb Pathog 152, 104658 (2021).

73. C. S. Chin et al., Nonhybrid, finished microbial genome assemblies from long-read SMRT sequencing data. Nat Methods 10, 563–569 (2013).

74. B. J. Walker et al., Pilon: an integrated tool for comprehensive microbial variant detection and genome assembly improvement. PLoS One 9, e112963 (2014).

75. B. J. Bennett et al., Trimethylamine-N-oxide, a metabolite associated with atherosclerosis, exhibits complex genetic and dietary regulation. Cell Metab 17, 49–60 (2013).

76. J. W. Jung et al., Mild hypercapnia before reperfusion reduces ischemia-reperfusion injury in hyperacute ischemic stroke rat model. J Cereb Blood Flow Metab, 271678X241296367 (2024).

77. J. H. Garcia, S. Wagner, K. F. Liu, X. J. Hu, Neurological deficit and extent of neuronal necrosis attributable to middle cerebral artery occlusion in rats. Statistical validation. Stroke 26, 627–634; discussion 635 (1995).

78. E. Z. Longa, P. R. Weinstein, S. Carlson, R. Cummins, Reversible middle cerebral artery occlusion without craniectomy in rats. Stroke 20, 84–91 (1989).

79. J. Chen et al., Intravenous administration of human umbilical cord blood reduces behavioral deficits after stroke in rats. Stroke 32, 2682–2688 (2001).

80. D. W. McBride, D. Klebe, J. Tang, J. H. Zhang, Correcting for Brain Swelling’s Effects on Infarct Volume Calculation After Middle Cerebral Artery Occlusion in Rats. Transl Stroke Res 6, 323–338 (2015).

81. K. Takada, A. Fukatsu, S. Otake, M. Hirasawa, Isolation and characterization of hemolysin activated by reductant from Prevotella intermedia. FEMS Immunol Med Microbiol 35, 43–47 (2003).

82. K. Lee et al., The ferrichrome receptor A as a new target for Pseudomonas aeruginosa virulence attenuation. FEMS Microbiol Lett 363, (2016).

83. R. P. Rennie, L. Turnbull, C. Brosnikoff, J. Cloke, First comprehensive evaluation of the M.I.C. evaluator device compared to Etest and CLSI reference dilution methods for antimicrobial susceptibility testing of clinical strains of anaerobes and other fastidious bacterial species. J Clin Microbiol 50, 1153–1157 (2012).

